# Cooperative Binding of Cytosolic Type III Secretion System Proteins to the Injectisome Revealed by Live-Cell Single-Molecule Localization Microscopy

**DOI:** 10.1101/2025.10.30.685657

**Authors:** Olivia I.C. de Cuba, Gopika A. Lekshmi, Caroline F. Tyndall, Andreas Gahlmann

## Abstract

Type III secretion systems (T3SSs) are employed by many Gram-negative bacteria to translocate virulent effector proteins into host cells. Secretion occurs through the T3SS injectisome, a multi-membrane-spanning biomolecular machine. While the overall structure of the injectisome is becoming increasingly well defined, the structural dynamics that regulate secretion remain unresolved. Particularly important for the functional regulation of type 3 secretion are the cytosolic injectisome proteins which transiently associate with injectisomes and whose structural dynamics and function remain incompletely understood. Here, we use long-exposure single-molecule localization microscopy to quantify the bound times of the cytosolic components SctQ, SctL, and SctN, at the injectisome of *Yersinia enterocolitica* both in controllably induced secretion ON and OFF states and in the presence and absence of effector proteins. In the absence of effector proteins, each component exhibits distinct short-lived binding behavior on a timescale of seconds. However, the presence of the effector protein YopE increases bound times and synchronizes binding behavior across components, specifically without active secretion. Upon activation of secretion, a subpopulation of long-lived binding events emerges for SctQ, SctL, and SctN even within the same injectisomes, indicating that binding behavior between these proteins is secretion state-dependent and cooperative. These findings establish that the interaction kinetics between cytosolic injectisome proteins are modulated by the presence of secretion substrates *and* secretion activating signals, and reveal a new layer of heterogeneity in the functional regulation of type 3 secretion.

## INTRODUCTION

Type III secretion systems (T3SSs) are conserved multi-membrane spanning protein complexes used by many Gram-negative pathogens to assemble either flagella or injectisomes that translocate effector proteins directly into eukaryotic cells^1^. The injectisome includes an extracellular needle connected through the bacterial envelope to a membrane-embedded export apparatus and a cytosolic protein complex that engages, energizes, and prioritizes effector proteins for secretion (Fig. 1a)^2–5^.

**Figure 1:**
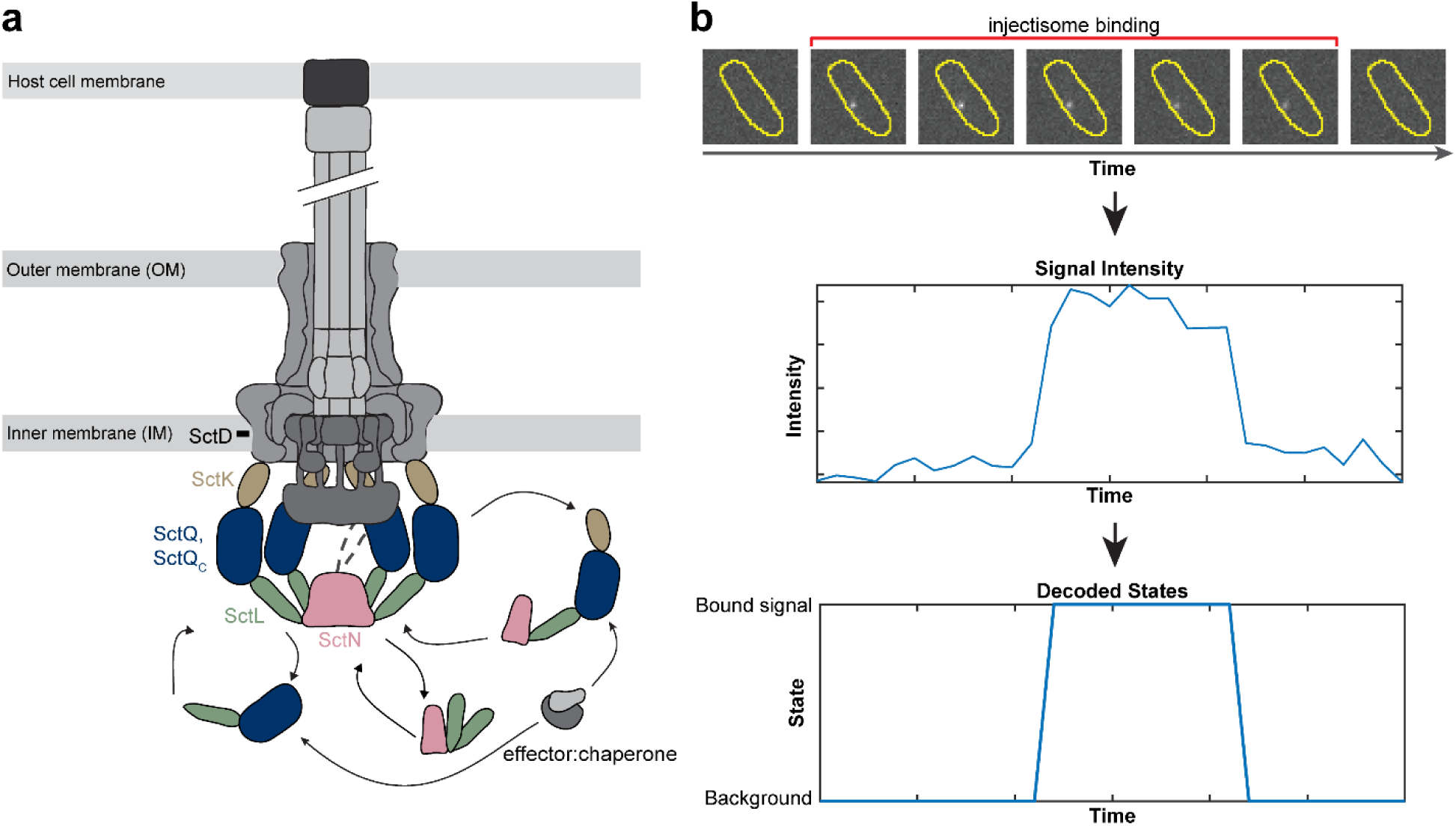
**(a)** Schematic of the Type III Secretion System (T3SS) architecture in *Yersinia enterocolitica*, highlighting the substructures and the cytosolic components analyzed in this study: SctQ, SctL, and SctN. The stoichiometry of these proteins at the injectisome have been estimated previously at 24 copies of SctQ^39^, 12 copies of SctL^39^, and 6 copies of SctN^40^. **(b)** Long-exposure imaging strategy used to extract bound times from stationary single-molecules. Injectisome-bound molecules appear as sharp foci, while cytosolic diffusing molecules blur into the background. A hidden-Markov model helps distinguish between the bound signal and background state, enabling an automated transition from signal intensities to decoded states used to measure bound times at individual injectisomes (see text for details).

The biogenesis of the T3SS follows a hierarchical process in which substrates are secreted in a defined order to assemble the functional injectisome and subsequently secrete effector proteins. Early substrates, such as the inner rod (SctI) and needle protein (SctF), are secreted first to assemble the needle structure^6,7^. This enables secretion of middle substrates, the translocator proteins (SctA, SctB, SctE), which form the translocon pore in the host membrane^8^. Once this pore is established, the system switches to secreting late substrates, the effector proteins, directly into the host cell. These transitions are controlled by molecular switches: the ruler protein SctP and export gate component SctU halt early substrate secretion once the needle reaches its final length, while the gatekeeper complex (SctW) dissociates from SctV to regulate the shift from translocator to effector secretion^9,10^. In *Yersinia enterocolitica*, an assembled injectisome can be reversibly turned ON and OFF using the calcium switch; high calcium represses secretion, while calcium depletion activates effector secretion^11–13^. While the cytosolic components play a role in the secretion of early, middle and late substrates, if and how the cytosolic components of the injectisome reorganize to accommodate these functional transitions is not yet fully understood.

*In situ* structural studies by cryo-electron tomography have converged on an architectural model in which the export apparatus is capped on its cytosolic face by a multiprotein assembly comprising SctK, SctQ, and SctL, arranged around the hexameric ATPase SctN and its central stalk (SctO)^2,3,14,15^. Initial *in situ* observations using *Shigella* and *Salmonella* minicell-producing mutants revealed six distinct cytoplasmic “pods” that tether SctN to the membrane-embedded SctD ring^2,16,17^. In contrast, in *Yersinia* a more continuous, ring-like density beneath the injectisome, reminiscent of the flagellar C ring, was observed^18,19^. These densities include SctK, SctQ, and likely SctQ_C_, the alternatively expressed C-terminal fragment of SctQ^20,21^, while SctL retains a six-fold symmetric spoke-like arrangement cradling SctN. Protein-protein interaction studies using bacterial hybrid systems^22–24^ and photo-crosslinking^25^ have mapped a sequential SctK–Q–L–N arrangement, with SctQ playing a particularly enigmatic role; its N-terminal domain has been implicated in interactions with SctK^25^, SctQ_C_^26^, SctN^23^, SctD^27^, and even chaperone-effector complexes^23,24,28^. Together, these findings provide useful information about which proteins interact at the injectisome, but they offer limited insight into how these proteins (re)organize in response to environmental cues.

Structural and mutational analyses have begun to define how SctQ, SctL, and SctN interact within this cytosolic assembly. Studies in *Salmonella* and *Shigella* demonstrate that the ATPase SctN uses a short N-terminal hook to bind a dimer of SctL, and that deletion of this N terminal domain disrupts both ATPase oligomerization and secretion^23,29^. SctL is organized into two functional regions, with a C-terminal segment (∼140-220) that interacts with SctN and an N-terminal adaptor domain (residues 1-30) that contacts C-terminal SctQ (SPOA) domains^22,26^. Within this N-terminal adaptor segment, a conserved hydrophobic triad mediates contacts with complementary grooves on the two C-terminal (SPOA) folds of SctQ^26^. In addition to these SPOA-mediated interactions, SctQ interacts with SctK through its N-terminal region^26^. Finally, electrostatic interactions between SctK and SctD dock the cytosolic complex proteins to the membrane^30^. SctD is an integral membrane ring protein with a transmembrane domain anchoring the injectisome and a cytosolic domain that interacts with SctK^31^. These interactions, supported by cryo-electron tomography and crosslinking studies, reinforce the sequential SctK–Q–L–N arrangement that bridges the export apparatus to the ATPase^2,25,32^. Mutations disrupting either the C-terminal (SPOA) interfaces of SctQ or the adaptor motif of SctL destabilize cytosolic complex assembly and abolish secretion^26,32^. These domains constitute the structural core architecture that underlies the (cooperative) binding kinetics examined in this study.

Complementing these structural insights, biochemical and genetic studies have illuminated a functional role for the cytosolic injectisome proteins in substrate secretion hierarchy. A high molecular weight complex comprising SctK, SctQ, and SctL was shown to coordinate the sequential loading of translocases (which form the translocon pore) and effector proteins, each guided by cognate chaperones^33^. Using native gel electrophoresis and subcellular fractionation in *Salmonella*, this study demonstrated that this so-called “sorting platform complex” remains intact even in the absence of the needle complex, suggesting it assembles independently and may serve as a pre-assembled substrate queue. Disruption of this sorting platform complex impairs secretion efficiency and abolishes substrate prioritization, underscoring the regulatory importance of this cytosolic interface^33^. Whether similar organizational principles apply across species remains to be determined. Together, these studies define the components and functional organization of the cytosolic interface, but they provide limited insight into how these proteins dynamically bind and unbind from the injectisome.

Fluorescence-recovery-after-photobleaching (FRAP) experiments in living cells have shown that the cytosolic interface of the injectisome is highly dynamic. In *Yersinia*, SctQ exchanges between injectisome-bound and cytosolic pools with a recovery half-time of roughly one minute, accelerating under secreting conditions and requiring ATPase activity, as recovery is abolished when ATPase function is (genetically) disrupted^34^. These data indicate that SctQ binding and unbinding is connected to secretion activity. Single-molecule tracking by our lab and others, in *Y. enterocolitica* subsequently revealed that SctQ, SctL, and SctN populate multiple diffusive states in addition to an injectisome-bound state^28,35,36^. These data are consistent with the formation of hetero-oligomeric complexes that assemble in the cytosol and remain stable for at least several hundreds of milliseconds and possibly longer. A recent study suggested that chaperones facilitate a cytosolic handover of effector proteins to cytosolic SctQ-containing complexes^37^. These SctQ- and effector-containing complexes were proposed to shuttle substrates to the injectisome, implying that substrate availability informs both the composition of cytosolic complexes and their binding to the injectisome^28,37,38^. Together, these observations suggest a dynamic picture in which cytosolic complex formation, substrate loading, and injectisome engagement are kinetically coupled.

Here, we use long-exposure single-molecule localization microscopy to extract the bound-time distributions (also referred to as residence times) of SctQ, SctL, and SctN at the injectisome of *Y. enterocolitica* in effector-free and YopE-expressing backgrounds as well as under secretion ON and OFF conditions. We also utilize 3D single-molecule tracking to quantify the bound fraction of SctQ as a proxy for the binding rate k_on_ under different secretion states and the presence/absence of YopE. This approach allows us to test whether secretion state and effector presence differentially regulate the binding kinetics and the number of resolvable bound state time constants. By quantifying kinetic parameters across defined environmental and substrate conditions, we provide new insights into how the T3SS cytosolic proteins dynamically adjust their interactions with injectisomes in response to secretion activating signals and secretion substrate availability.

## RESULTS

To quantify the bound times of cytosolic T3SS components at the injectisome, we used a biosafety level 1-compatible *Yersinia enterocolitica* strain lacking six major effector proteins referred to as T3SS^Δeffector^ (also known as IML421asd, see Methods, list of strains)^4^. We expressed Halo-tagged SctQ and SctL from their native loci via allelic exchange; Halo-SctN was expressed from a plasmid due to unsuccessful allelic replacement attempts. All fusion proteins were confirmed to be stable (Fig. S1a) and functional (Fig S1b), i.e. capable of reversible induction of secretion by the calcium effect (Fig S1c). These strains thus allowed us to probe the binding behavior of SctQ, SctL, and SctN in the absence of effector proteins.

A small fraction of Halo-fusion proteins (1-5 per cell) was visualized using low concentrations of dye coupled with long-exposure time imaging. Under these conditions, fast-moving molecules blur into the background, and stationary, injectisome-bound proteins are detected as sharp point spread functions (Fig 1b). The pooled bound times were displayed as probability density function (pdf)-normalized histograms and fit with single- or bi-exponential decays. The resulting bound time constant, τ, is the inverse of the dissociation rate of the respective protein from the injectisome, k_off_.

### Effector-Free Conditions Reveal Distinct Binding Dynamics for Injectisome-Associated Proteins SctQ, SctL, and SctN

In the T3SS^Δeffector/OFF^ condition, the bound time histograms of SctQ, SctL and SctN are best fit by single exponential decays (Fig. 2a-c). Notably, all three proteins exhibit bound times on the order of a few seconds, but the fitted bound time constants are all distinct. SctQ exhibits the shortest bound time constant, τ = 2.5 s [2.4, 2.5], while the bound times constant of SctL and SctN are slightly longer, τ = 3.4 s [3.3, 3.4] and τ = 3.0 s [2.9, 3.0], respectively (the square brackets represent the 99% confidence intervals).

**Figure 2:**
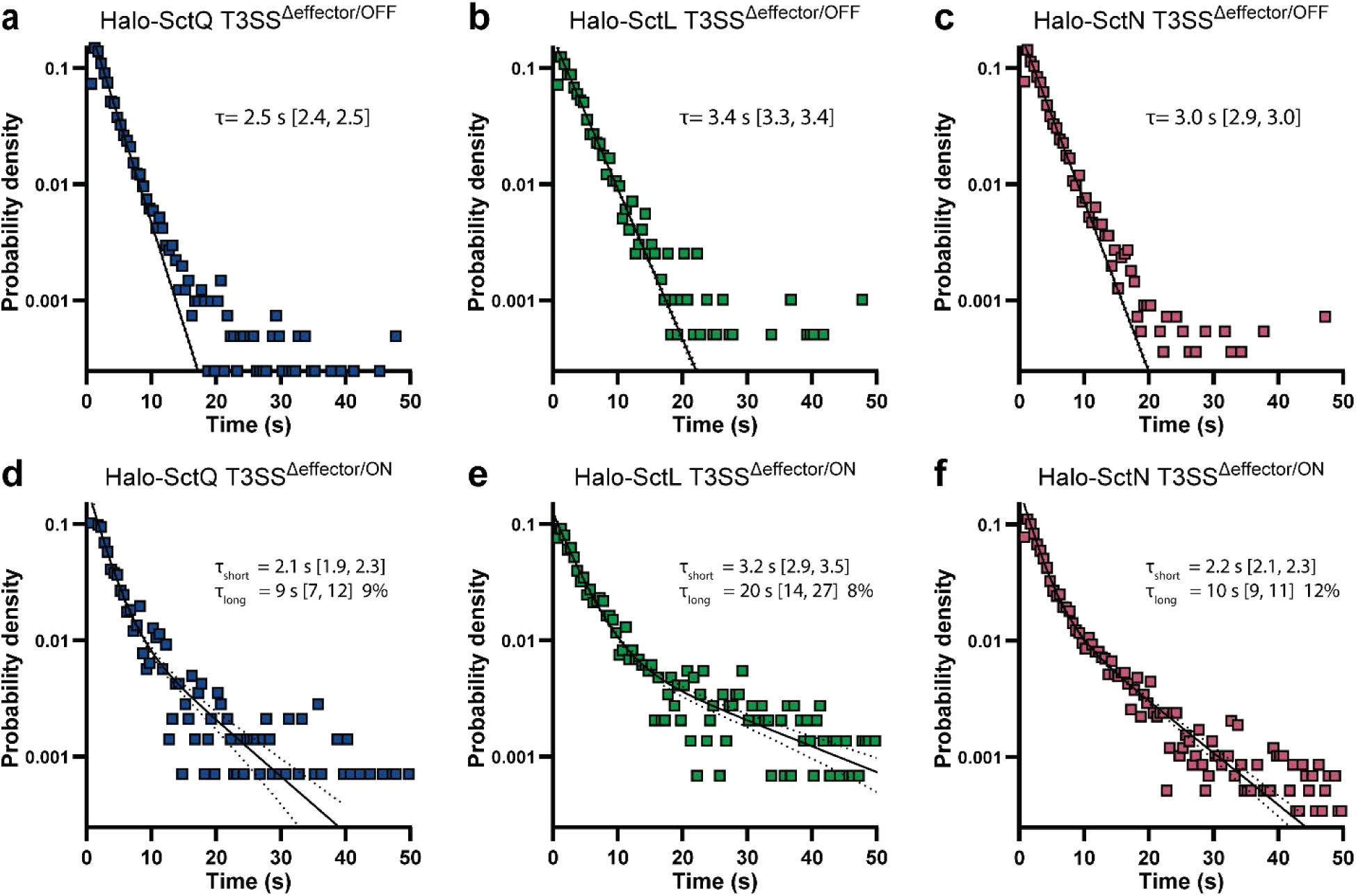
Distinct bound times for SctQ, SctL, and SctN and emergence of long-lived binding events under secretion ON conditions. Bound time distributions for Halo-tagged SctQ, SctL, and SctN under secretion OFF (top row) and ON (bottom row) conditions in the effector-free background. **(a–c)** Under secretion OFF conditions, all three proteins exhibit mono-exponential decay kinetics with distinct bound times: SctQ (τ = 2.5 s), SctL (τ = 3.4 s), and SctN (τ = 3.0 s). **(d–f)** Under secretion ON conditions, a subpopulation of long-lived binding events emerges for all proteins, requiring bi-exponential fits. Long-lived events comprise 9% (SctQ), 8% (SctL), and 12% (SctN) of trajectories, indicating two binding states at the injectisome. The total counts for the histograms in a-f are *n* = 4063, *n* = 1973, *n* = 5545, *n* = 1417, *n* = 1463, and *n* = 5857, respectively.

Under secretion ON conditions, we observed a stark change in the binding behavior of SctQ, SctL, and SctN at the injectisome. Rather than a simple difference in bound time constant, the single-molecule bound time data revealed a subpopulation of binding events with substantially longer durations (>10 s), necessitating bi-exponential fits to the bound time distributions. These long-lived events comprised approximately 9% of SctQ, 8% of SctL, and 12% of SctN trajectories (Fig. 2d-f), indicating the emergence of a distinct and more long-lived binding mode at the injectisome. The finding of long-lived states across all three proteins points to a coordinated and potentially cooperative injectisome-bound configuration under secretion ON conditions.

Notably, the short component of the bi-exponential fit was shorter under ON than OFF conditions: for example, SctQ’s short-term bound time decreased from 2.5 s (OFF) to 2.1 s (ON). When considering both short- and long-lived events, the weighted average bound time constants were 2.7 s for SctQ, 4.4 s for SctL, and 3.1 s for SctN under T3SS^Δeffector/ON^ conditions. Thus, despite the prevalence of shorter short-lived events in our single-molecule data, the bound times constant did not skew toward faster exchange overall under secreting conditions, as previously reported using FRAP^34^. Because FRAP half-times average over relatively few individual injectisomes, we asked whether individual injectisomes engage in both short- and long-lived binding, or whether distinct classes of injectisomes exhibit either short or long-lived binding modes only.

### The Same Injectisome Can Engage in Short and Long Binding Modes

Within the same bacterial cell, we observed both short- and long-lived binding events for SctQ, SctL, and SctN (Fig. 3a), establishing that the emergence of long-lived states is not due to cell-to-cell variation. This observation is possible because each single-molecule binding event can be localized to a specific cell and even to a specific subcellular location (Fig. 1b) and repeated binding events at the same spatial locations, i.e. clusters of single-molecule localizations, can be used to approximate the position of injectisomes. To assess whether long-lived binding events preferentially occur at a distinct subset of injectisomes, we clustered binding events within each field of view using DBSCAN (ε = 150 nm, minPts = 2) and classified events as long if their bound time exceeded 15 seconds. For injectisomes with multiple binding events, we found subsets showing exclusively short-lived events, exclusively long-lived events, as well as mixed duration events (Fig. 3b, table S2).

**Figure 3:**
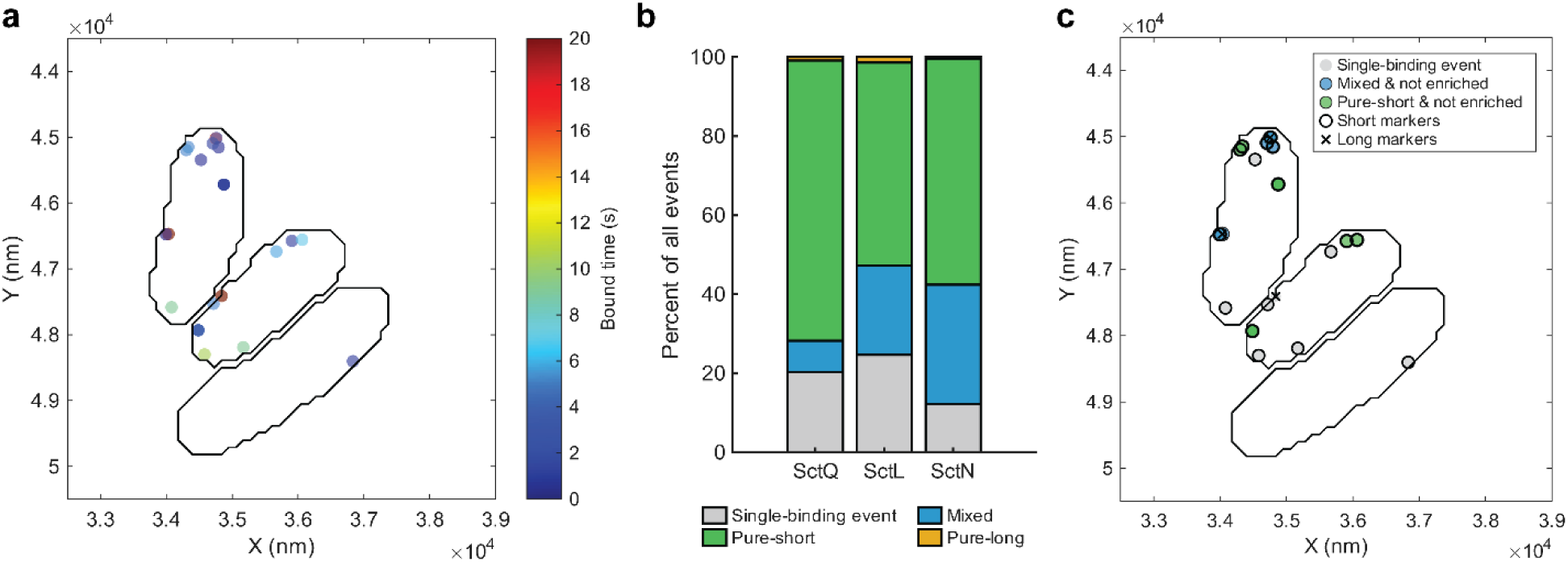
Short- and long-lived binding events occur at the same injectisome. **(a)** Representative cells showing spatial co-localization of short- and long-lived binding events for SctQ, SctL, and SctN, confirming that both binding modes occur within the same cell and at the same subcellular locations. **(b)** Classification of injectisomes based on binding profiles: mixed (both short and long), short-only, and long-only. (Breakdown of all counts analyzed can be found in Table S2.) **(c)** Statistical analysis using a hypergeometric test with FDR correction (ε = 100 nm, minPts = 2; long = dwell ≥ 30 frames) reveals no significant enrichment of long-lived events at specific clusters, indicating that binding mode heterogeneity is not spatially segregated.

We note that these data were obtained by sparsely labeling the proteins of interest. The abovementioned injectisome classification could therefore be skewed because we are not sampling all the possible binding events at a given injectisome. We therefore compared the observed fraction of long events within clusters to the global fraction of long events across the field of view. Using a hypergeometric test with Benjamini–Hochberg correction (α = 0.05), we asked whether clusters contained more long-bound events than expected under random distribution. No clusters showed significant enrichment of long events for any protein, indicating that long-bound events were not spatially localized beyond random expectation. We repeated this analysis for short-bound events and again observed no statistical evidence for enrichment. Based on this analysis, we conclude that long- and short-lived bound states are not spatially segregated and occur at the same injectisome during the 22-min time windows of data acquisition.

### YopE Modulates Injectisome Binding Dynamics, With SctQ Showing Distinct Sensitivity Across Secretion States

To determine how the presence of effector proteins influences injectisome-associated protein dynamics, we quantified the bound lifetimes of SctQ, SctL, and SctN under secretion ON and OFF conditions in the presence of the effector protein YopE. YopE was constitutively expressed from a plasmid and confirmed to be successfully secreted in a Ca^2+^-dependent manner in all strains (Fig S1b). (SycE, the cognate chaperone of YopE is natively expressed from the pYV virulence plasmid in all of the strains used in this study.)

In the presence of YopE, all three proteins exhibited increased bound times compared to the effector-free condition (T3SS^Δeffector/OFF^), and their bound time constants converged to statistically indistinguishable values (Fig. 4a-c). Mono-exponential fits yielded bound time constants of τ = 4.1 s [4.0, 4.2] for SctQ, τ = 3.9 s [3.8, 4.0] for SctL, and τ = 3.9 s [3.8, 3.9] for SctN. The increase was most pronounced for SctQ, which nearly doubled its bound time constant relative to the effector-free condition. These findings indicate that effector presence alone can increase the bound time of cytosolic components at the injectisome in secretion OFF conditions.

**Figure 4:**
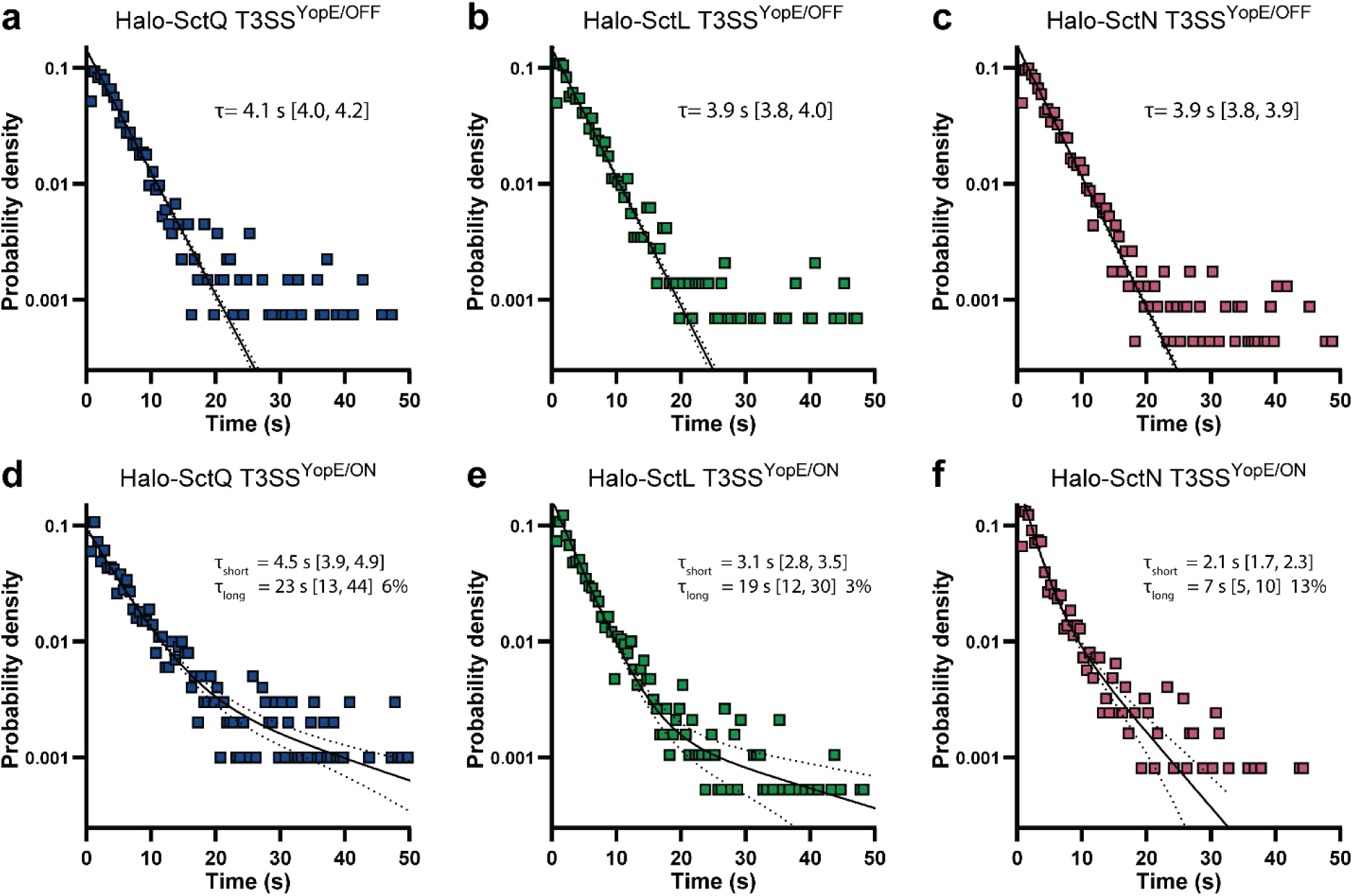
Effector presence increases bound times and synchronizes binding behavior. Bound time distributions for SctQ, SctL, and SctN under secretion OFF (top row) and ON (bottom row) conditions in the presence of the effector protein YopE. **(a–c)** Under secretion OFF conditions, all three proteins show increased bound times compared to the effector-free background, with mono-exponential fits yielding τ ≈ 4 s for all proteins. **(d–f)** Under secretion ON conditions, SctQ exhibits both an increased short bound time (τ = 4.5 s) and a distinct long-lived subpopulation (τ = 23 s, 6% of events). SctL and SctN show smaller changes, indicating component-specific sensitivity to effector presence. The total counts for the histograms in a-f are *n* = 1345, *n* = 1446, *n* = 2292, *n* = 1003, *n* = 1902, and *n* = 1240, respectively.

Next, we asked whether these effects persist during active secretion (T3SS^YopE/ON^). As in T3SS^Δeffector/ON^, the T3SS^YopE/ON^ condition revealed a subpopulation of binding events with substantially longer durations (>10 s), again prompting bi-exponential fits to the bound time distributions. These long-lived events comprised 6% of SctQ, 3% of SctL, and 13% of SctN trajectories (Fig. 4d-f). As before, we observed mixed long- and short-bound behavior at single injectisomes (Table S3) and there was no statistical evidence for enrichment of any one type of binding at specific spatial locations (Fig S3), again ruling out the possibility of two distinct classes of injectisomes. SctQ exhibits the longest short-bound time (τ = 4.5 s [3.9, 4.9]), while the short-bound times of SctL and SctN are τ = 3.1 s [2.8, 3.5] and τ = 2.1 s [1.7, 2.3], respectively. Thus, the presence of the YopE effector more than doubles the short-bound duration for SctQ, while leaving the short-bound durations for SctL and SctN largely unchanged.

### SctQ Injectisome-Bound Fraction Is Not Sensitive to Secretion State or YopE Effector **Presence**

The observation that SctQ exhibits slightly longer bound time constants in the secretion ON state does not match previous FRAP measurements that indicate faster protein exchange under secreting conditions^34^. Protein exchange at injectisomes is a function of both the binding rate constant (k_on_) and the unbinding rate constant (k_off_). The above bound time measurements only report on the k_off_ and so do not rule out changes in the association rate constant. An increase in k_on_ would increase the fraction of injectisome bound molecules, especially if bound time remains constant or increases. However, all available binding sites at the injectisome may already be occupied or rapidly reoccupied under saturating conditions, in which case we expect bound fraction to remain constant across our experimental conditions.

Our previous 3D single-molecule tracking results in *Y. enterocolitica* under secretion ON conditions found freely-diffusing cytosolic states for eYFP-SctQ in addition to a prominent stationary state corresponding to injectisome-bound SctQ^35,36^. Similar results are obtained for Halo-SctQ (Fig. 5a, left panel), under all conditions described above. The fraction of slow-moving SctQ trajectories (mean displacement < 4 nm/ms) thus enables us to estimate the SctQ bound fraction (Fig. 5a, right panel), which may be sensitive to changes in the association rate constant k_on_. Quantifying the injectisome bound fraction of SctQ using this metric showed very small differences across conditions.

**Figure 5:**
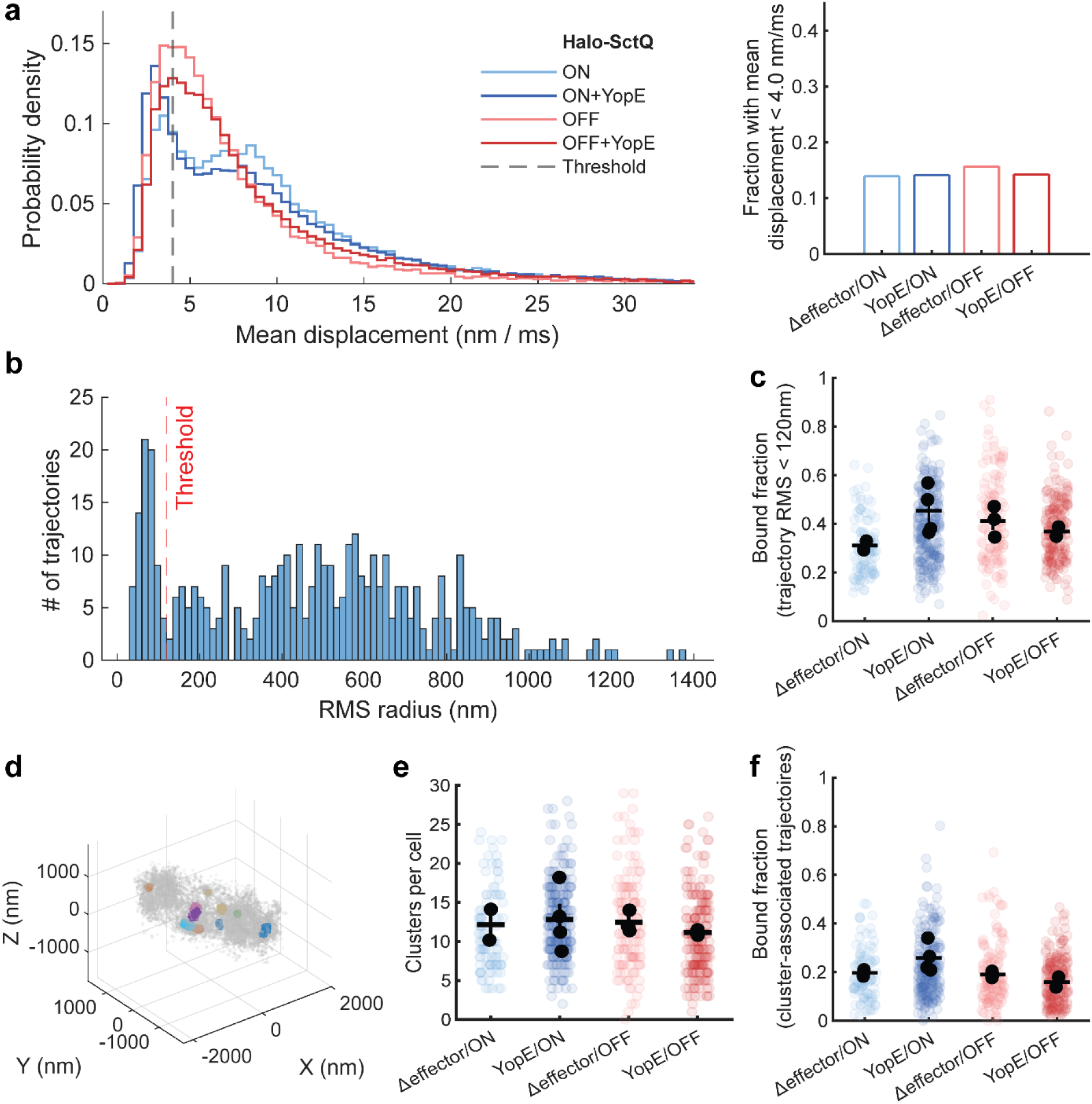
SctQ injectisome-bound fraction is not sensitive to secretion state or YopE effector presence. **(a, left)** Pooled single-molecule displacements of Halo-SctQ under four experimental conditions. For each condition, at least 100 cells and 10 000 trajectories were analyzed (see table S4 for exact counts). **(a,** right) Quantification of the fraction of slow-moving trajectories (mean displacement < 4 nm/ms) per cell, used as a proxy for injectisome association. **(b)** Histogram of single-molecule trajectory RMS deviations, representative cell, threshold at 120 nm was selected based on tall peak to the left of that threshold, which appears robustly for all cells analyzed. **(c)** RMS threshold-based analysis confirms consistent injectisome-bound fraction across conditions, with minor variation that follows the same patterns as the mean number of injectisomes (see panel e), supporting the conclusion that injectisome bound fraction stays constant across conditions. **(d)** Representative localizations of Halo-SctQ with identified localization clusters: a proxy for injectisomes. **(e)** Number of clusters identified per cell by the clustering analysis of DBSCAN within 60 nm (see methods). Localization clusters are consistent with (repeated) binding to a spatially confined region, i.e. injectisome. The finding of ∼12 injectisomes per cell is consistent with previous cellular injectisome counts by other methods^28,41,42^. **(f)** Cluster-based categorization of single-molecule trajectories confirms consistent injectisome-bound fractions across conditions. **(c, e, f)** Bars indicate the mean across biological samples, with error bars representing the standard error of the mean across samples. Colored points show individual cells, while black points denote per-sample means.

To alternatively assess potential variations in injectisome association, we implemented an approach based on the root mean squared (RMS) radius, also called the radius of gyration. This quantifies how spatially spread out the trajectory is around its mean position. For a molecule associated with an injectisome, we expect the trajectory to be spatially confined, with the apparent motion being predominantly due to localization error. When plotting the RMS radius spread for trajectories, we consistently observed a peak below 120 nm and selected this to be the upper threshold for injectisome association (representative cell shown in Fig. 5b). By classifying trajectories below the threshold as injectisome-associated, a bound fraction was computed for the different conditions (Fig. 5c). Here too, we found only small variations across our conditions with a series of unpaired t-tests indicated no statistically significant differences (table S5).

As a third approach, we used spatial clustering of single-molecule localizations. By identifying localization clusters (at the membrane), consistent with injectisome locations (Fig. 5d)^35,36^, and classifying single-molecule trajectories as cluster-associated or not, we obtained yet another measure of the injectisome bound fraction (Fig. 5f). Similar to the previous approach, this analysis also revealed that the fraction of cluster-associated trajectories did not change significantly across our experimental conditions, as any observed differences were within statistically expected variation (table S6).

Taken together, these data show that the injectisome-bound fraction of SctQ does not vary with secretion state or effector presence. We therefore focus our discussion on the observed changes in injectisome bound times only.

## DISCUSSION

During infection, *Yersinia enterocolitica* must rapidly adapt to changing host environments. A rapid transition between the secretion OFF and ON state is thought to be essential for evading immune clearance and thereby persistence at sites of lymphoid colonization^43–47^. Activation of secretion, as well as deactivation, can occur rapidly^47,48^ in response to external stimuli, suggesting strong external sensing mechanisms that modulate secretion, possibly through the cytosolic injectisome components.

In this study, we measured the bound times of SctQ, SctL, and SctN under secretion OFF and ON conditions in the presence and absence of the effector protein YopE. In secretion OFF conditions, we observed mono-exponential decay kinetics for SctQ, SctL, and SctN, indicating a single, transient bound state at the injectisome for each protein. In the absence of effectors (T3SS^Δeffector^), each protein exhibited a distinct fitted bound time, with SctQ having the shortest, followed by SctN and SctL, suggesting that these components do not both bind *and* unbind as an assembled unit. Independent exchange of these proteins is a notion additionally supported by a recent study showing that SctQ can still bind to and dissociate from the injectisome even when the inner membrane ring protein SctD is crosslinked and thus unable to exchange itself^49^.

In contrast, the fitted bound times of all three proteins converged to approximately 4 seconds in the presence of the effector protein YopE in the secretion OFF state. This synchronization of bound times shows that effector presence alone, independent of active secretion, can modulate the binding kinetics of cytosolic components. Moreover, this convergence of bound times is consistent with the notion that SctQ, SctL, and SctN both bind and unbind as a part of the same protein complex and is consistent with effector proteins that are preloaded onto cytosolic SctQ-L-N complexes^36,50^ that then subsequently bind to injectisomes^37^. Such a mechanism could help prime the system for rapid activation of secretion. The convergence of bound times may reflect a shift toward a more uniform or stabilized interaction mode of SctQ, SctL, and SctN at injectisomes, perhaps due to effector-induced conformational or compositional remodeling. An alternative explanation to intact complex dissociation would be that all other proteins disassemble immediately after at least one of them unbinds from the injectisome. Such a model is supported by gene deletion studies that have established that cytosolic injectisome proteins are mutually dependent on each other injectisome binding^2,32,34,51^, expression of each protein is required for injectisome assembly^2,16,52^ and type 3 secretion^22–24,29,53^. Regardless of the exact mechanism at play, the continued presence of unsecreted YopE effectors seems to induce a coordinated, bind-and-reject process, where a SctQ-L-N complex binds and unbinds together. Consistent with this interpretation, our data show that such temporally coordinated binding and unbinding indeed disappears in the secretion ON state, in which YopE is secreted and thereby removed from injectisomes.

Under secretion ON conditions and irrespective of whether YopE is present, SctQ, SctL, and SctN all display a subpopulation of long-lived binding events, resulting in bi-exponential decay kinetics. Detection of this long-lived bound state is uniquely enabled by single-molecule measurements, which allow direct quantification of injectisome bound times of *individual* protein molecules. Importantly, we observed both short- and long-lived binding events at the same subcellular locations, suggesting that individual injectisomes can engage proteins in different binding modes, one mode being more stable than the other. This raises the intriguing possibility of functional asymmetry within single injectisomes, either through structural heterogeneity within an injectisome or through dynamic switching between distinct operational states in time (our pooled single-molecule data were obtained over a timescale of ∼20 minutes). In support of the structural heterogeneity hypothesis, cryoEM studies have shown that the ATPase SctN forms an asymmetric hexamer at the base of the injectisome^54^, raising the possibility that asymmetry extends to other cytosolic injectisome components and contributes to the observed kinetic heterogeneity. Future structural and computational studies will be essential to determine whether the short- and long-bound states correspond to distinct conformational or functional states of the injectisome under defined secretion conditions.

Among the three cytosolic components analyzed, SctQ consistently exhibited the most pronounced changes in bound time across conditions and was particularly sensitive to the presence of the effector protein YopE. Under secretion OFF conditions, the addition of YopE nearly doubled the SctQ bound time, bringing it into close alignment with SctL and SctN. Under secretion ON conditions, SctQ again showed heightened sensitivity, with both an increased short-bound time and a subpopulation that fits to a longer-bound time constant. This component-specific response supports the notion that SctQ serves in a key regulatory role within the assembled injectisome.

Previous studies have examined complete turnover of the SctQ subunits at the injectisome under secretion ON conditions similar to those reported here, as well as a secretion OFF state prior to injectisome assembly^34,49^ (which is unlike that reported here). We can also interpret our single molecule bound state lifetimes in the context of complete ring turnover. Assuming 24 SctQ subunits^39^ independently exchanging at the observed mixture of lifetimes, the expected time for *all* sites at the injectisome to undergo at least one unbinding event is approximately 21 s under T3SS^YopE/ON^ conditions and 10 s T3SS^Δeffector/ON^ (see SI for further explanation). Similarly, we estimate approximately 16 s under T3SS^YopE/OFF^ and 10 s for T3SS^Δeffector/OFF^. FRAP-based measurements of subunit exchange are sensitive to both k_on_ and k_off_, but are also influenced by additional factors such as photobleaching and binding site occupancy (see SI), which can slow the apparent turnover rate. Taken together, our single-molecule results are qualitatively consistent with previous FRAP observations under similar secretion ON conditions^34,49^.

Furthermore, our results are compatible with a shuttling model in which SctQ-containing complexes bind effectors, such as YopE, in the cytosol and then deliver the effectors to the injectisome for secretion^37^. We evaluated whether such a mechanism alone could support the total known per-cell secretion rate^55^ of 7–60 molecules s⁻¹. Assuming near-saturated occupancy of injectisome binding sites, the fast dissociation rate (derived from τ_short_) sets an upper bound on secretion throughput. Using the experimentally measured lifetime of Halo–SctQ under T3SS^YopE/ON^ conditions (τ ≈ 4.5 s) and an average of 12 injectisomes per cell (Fig 5e), predicted secretion rates can reach up to 64 molecules s⁻¹(see SI for further explanation). Previously reported secretion fluxes are 7–60 molecules s⁻¹, indicating that SctQ shuttling and turnover alone could support effector export even if not every binding event results in a secretion event. We note however that while our results show injectisome-associated dynamics on a timescale compatible with effector translocation, they do not provide direct evidence for an effector shuttling mechanism. Indeed, direct binding of effectors to the injectisome is an alternative and not mutually exclusive mechanism. Future work may be able to distinguish between these mechanisms.

In conclusion, our analysis of injectisome-bound lifetimes reveals that cytosolic components exhibit secretion state- and effector-dependent binding modes. In secretion OFF states, distinct bound times for SctQ, SctL, and SctN indicate independent binding and/or unbinding. However, the presence of YopE synchronizes these bound states, consistent with cooperative binding within a SctQ–L–N–YopE complex engaging in a coordinated “bind-and-reject” cycle. Under secretion ON conditions, YopE is secreted and this temporal coordination disappears: biexponential bound time distributions arises as short- and long-bound states manifest at the same injectisome for SctQ, SctL, and SctN. Importantly, comparing effector-free and YopE-expressing strains allowed us to separate the effects of secretion state from those caused by the effector YopE. The fact that cytosolic injectisome protein binding kinetics are altered even in the absence of active secretion supports the idea that cytosolic remodeling, potentially through effector or effector–chaperone interactions, can precondition cytosolic injectisome proteins before injectisome association, and alter their binding behavior at the injectisome. By directly linking effector presence and secretion state to injectisome-bound lifetimes, we show that type 3 secretion does not arise from a static injectisome architecture, but from dynamically changing, cooperative binding kinetics among cytosolic injectisome proteins.

## METHODS

### Bacterial culture

*Yersinia enterocolitica* strains were grown overnight in Brain Heart Infusion (BHI, Sigma-Aldrich, St. Louis, MO) media supplemented with nalidixic acid (NaI) (35 μg/mL) and 2,6-diaminopimelic acid (DAP) (80 μg/mL) at 28°C with shaking. Additional antibiotics were included when appropriate: kanamycin (50 µg/mL) and/or chloramphenicol (25 µg/mL). On the day of imaging, overnight culture was back diluted to a target OD_600_ of 0.1-0.3 and grown at 28°C with shaking for 1-1.5 h. For secreting conditions, glycerol (4 mg/mL), MgCl_2_ (20 mM), and EGTA (5 mM) were then added to the culture, and the culture was transferred to a 37°C water bath or incubator with shaking, for 3h to induce expression of the *yop* regulon and ensure secretion activation. For non-secreting conditions (post-secretion), cells were pelleted by centrifugation at 5,000 × *g* for 3 min and resuspended with fresh BHI supplemented with NaI, DAP, CaCl_2_ (5 mM) and grown for an additional hour.

### Strains and Plasmids

Plasmids were constructed using Gibson Assembly. Briefly, DNA fragments with overlapping ends were generated by Q5 PCR (New England Biolabs (NEB), Ipswich, MA) and assembled in a single isothermal reaction. Assembled plasmids were transformed into chemically competent *dh5α E coli* (NEB) and plated on LB agar supplemented with the appropriate antibiotic for selection. Individual colonies were cultured overnight, and plasmid DNA was isolated using a commercial miniprep kit (Qiagen, Hilden, Germany). Plasmid sequences were verified by long-read sequencing (Plasmidsaurus, Eugene, OR). Plasmids were transformed into the appropriate *Yersinia Enterocolitica* strain by electroporation.

List of oligonucleotides

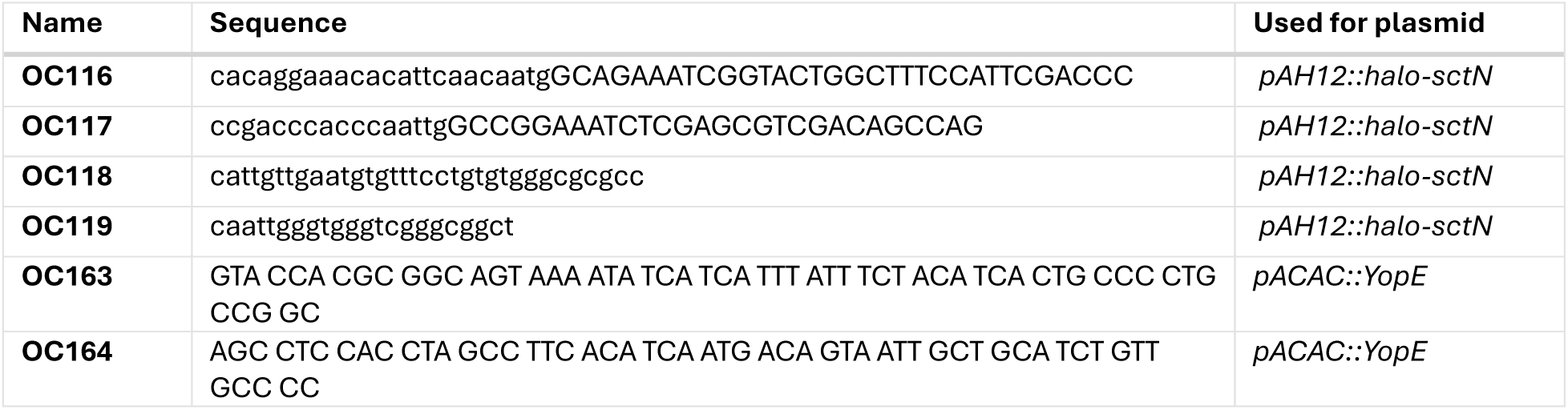

List of plasmids

| Plasmid | Genotype | Reference |
| --- | --- | --- |
| **1** | *pAH12::halo-sctN* | this study |
| **2** | *pAH12::halo* | this study |
| **3** | *pAH12::mEYFP* | this study |
| **4** | *pACAC::YopE* | this study |

List of Strains

| Strain | Genotype | Reference |
| --- | --- | --- |
| IML421asd<br>(ΔHOPEM<sub>T</sub>asd) | MRS40 *yopO*Δ12-427 *yopE*21 *yopH*Δ11-352 *yopM*23 *yopP*23 *yopT*135 Δ*asd* | 4 |
| AD4461 | IML421*asd halo-sctQ* | 32 |
| ADTM4525 | IML421*asd halo-sctL* | 32 |
| AD4165 | IML421*asd ΔsctN* | 34 |

### Secretion Assay

Cultures were grown as above, to desired secretion state. A small volume of culture was taken to measure the OD_600_. Main culture was pelleted (20,000 × *g*, 10 min, 4°C) and syringe filtered with a 0.22 µm PVDF filter (Millipore, Burlington, MA). Supernatant was mixed with trichloroacetic acid (TCA) to a final concentration of 10% w/v and incubated overnight at 4°C to precipitate proteins within the supernatant. Proteins were collected (20,000 × *g*, 20 min, 4°C) and washed with cold acetone (20,000 × *g*, 20 min, 4°C). The pellet was dried at room temperature and resuspended in reducing SDS-PAGE loading buffer (SDS [3%, wt/vol], glycerol [7.5%, wt/vol], 1.5% beta-mercaptoethanol, 0.0125% w/v Coomassie blue, buffered with Tris [37.5 mM]; pH 7.0). Loading volume was normalized across samples ran on the same gel, and proteins were separated by SDS-PAGE on 4–20% polyacrylamide gels (Bio-rad, Hercules, CA). For visualization, the gels were stained with Coomassie Blue and imaged with Bio-rad Gel Doc EZ imager.

### In-gel fluorescence Assay

Cultures were grown as above, to desired secretion state. HaloTag fusions were incubated with JFX-554 (Promega, Madison, WI) for the last 30 min of culturing. A small volume of culture was taken to measure the OD_600_. Culture was pelleted (5,000 x *g*, 5 min) and lysed in a buffer of 20 mM Tris HCl, 50 mM NaCl, pH 7.4, supplemented with 0.2 mg/mL of lysozyme and cOmplete™, Mini, EDTA-free Protease Inhibitor Cocktail (Roche, Basel, Switzerland). Samples were resuspended in SDS-PAGE loading buffer and heated for 5 minutes at 98°C before loading. Loading volume was normalized across samples ran on the same gel, and proteins were separated by SDS-PAGE on 4–20% polyacrylamide gels (Bio-rad). For visualization of in-gel JFX-554 fluorescence, a Biorad ChemiDoc MP system was used.

### Cell Plating for Microscopy

Cultures were grown as described above. In the case of HaloTag visualization, Janelia Fluor HaloTag ligand dye, JFX-554 (Promega) was added to the culture 30 minutes prior to imaging. The appropriate dye concentration was strain dependent: 0.5 nM for Halo-SctQ, 2 nM for Halo-SctL, 2nM for Halo-SctN. Cells were pelleted by centrifugation at 5,000 × *g* for 3 min and washed three times in fresh M9 minimal media. After the three washes, the remaining pellet was resuspended in M9 supplemented with DAP, MgCl_2_, glycerol, and EGTA (secreting conditions) or CaCl_2_. For super-resolution fluorescent imaging, fluorescent fiducial markers (Invitrogen) were added to cell suspensions. Cells were finally plated on 1.5% agarose pads in M9 containing DAP and EGTA or CaCl_2_.

### Optical Setup

Image data were acquired on a custom-built inverted fluorescence microscope based on the RM21 platform (Mad City Labs, Madison, WI), as previously described^35,56^. Immersion oil was placed between the lens objective (UPLSAPO 100; numerical aperture [NA], 1.4) and the glass coverslip (VWR, Radnor, PA; number 1.5; 22 mm by 22 mm). A 561 nm laser (Coherent, Santa Clara, CA; Genesis MX561-4000 MTM) was used for excitation of JFX-554 (Promega Madison, WI) (∼600 mW cm^−2^ for single molecule imaging at 25 ms exposure). Zero-order quarter-wave plates (Thorlabs, Newton, NJ, WPQ05M-561) were used to circularly polarize the excitation laser. A dichroic mirror (Chroma zt440/488/561rpc) was used to separate excitation/activation light from fluorescence emission of JFX-554.

Fluorescence emission was passed through a filter set including long-pass (514nm), short-pass (800 nm), and notch filter 561 nm/10bp (Semrock, Rochester, NY; LP02-514RU-25, Chroma, Bellows Falls, VT; ET700SP-2P8, and Chroma, Bellows Falls, VT; ZET561NF, respectively). In the case of 3D super-resolution imaging, the emission path contains a wavelength-specific dielectric phase mask (Double Helix, LLC, Boulder, CO) that is placed in the Fourier plane of the microscope to generate a double-helix point spread function (DHPSF)^57,58^. The fluorescence signal is detected on a scientific complementary metal oxide semiconductor (sCMOS) camera (Hamamatsu, Bridgewater, NJ; ORCA-Flash 4.0 V2). A flip mirror in the emission pathway enables toggling the microscope between fluorescence imaging and phase-contrast imaging modes without having to change the lens objective of the microscope. Illumination for phase contrast imaging was achieved with an LED at 625 nm (Thorlabs, Newton NJ; M625L3).

For long-exposure imaging, the laser intensity was attenuated significantly. Up to 2500 frames were collected per field-of-view with an exposure time of 500 ms.

### Long Exposure Data processing

Broadly, image processing occurred in a two-step process. First, potential binding events were identified using ThunderSTORM^59^ generating regions of interest for further analysis. Second, intensity traces over time were fitted with a Hidden Markov Model to distinguish between a bound and background state.

Specifically, prior to PSF fitting, images from each field-of-view were background subtracted and cropped as needed. PSFs were fit using ThunderSTORM in ImageJ using the difference of Gaussians (DoG) filter option with a threshold of 1.5*std(Wave.F1)^59^. Candidate binding events were further narrowed down with a requirement for there to be a localization within 50 nm at least 2 times per 1000 frames. The image stack and its corresponding ThunderSTORM data was imported into MATLAB. Images were normalized and converted to photoelectron units. A simple nearest-neighbor linker associated detections across frames (distance threshold 300 nm; maximum frame gap 15) to form candidate bound trajectories.

For every frame within a trajectory, signal was quantified as the sum in a 5×5 pixel ROI centered on the trajectory position. Pooled photon values were modeled with a Gaussian mixture model (GMM; 3 components) to estimate background and signal intensity distributions. Per-track ROI time series were then segmented with a two-state hidden Markov model (HMM; background ↔ signal): initial state means/variances were set from the GMM, a conservative transition matrix was used and the most probable state sequence was decoded by Viterbi. Quality gates rejected traces with (i) peak intensity ≤ background upper bound, (ii) minimum intensity ≥ signal lower bound, (iii) dynamic range below the GMM-implied separation, or (iv) substantial signal/background standard-deviation overlap. Error codes were logged for each exclusion. In addition, any bound state event longer than 40 frames (20 seconds) was inspected and manually approved. Remaining signal segments were taken as valid dwell events; durations (frames) were converted to time using 0.5 s/frame and compiled across tracks.

### Determination of t_bound_

To characterize the distribution of bound state lifetimes, dwell time measurements were extracted for each condition and protein variant and binned into normalized histograms. To reduce the influence of short-duration events on the fit, the first histogram bin (0-0.5 seconds), a group that is disproportionally undercounted, was excluded from model fitting. For each distribution, both single- and double-exponential decay models were fitted to the data using nonlinear least-squares regression. The single-exponential model was defined as y(t) = A e^−λ^ ^t^ and the double-exponential model as *y*(*t*) = *A*_1_*e*^−λ1*t*^ + *A*_2_*e*^−λ2*t*^ with time constants 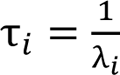.

Model selection was based on comparison of the Akaike Information Criterion (AIC), with the bi-exponential model favored if AIC was reduced by more than 2 units. In addition, if a state in the bi-exponential fit contributed less than 2% of the population, the mono-exponential model was favored. Goodness-of-fit was assessed via the coefficient of determination (R²). For each fit, 99% confidence intervals were estimated by propagation of uncertainty using the Jacobian and parameter covariance matrix. Fits and their confidence intervals were overlaid on probability density plots to visualize lifetime distributions for each experimental condition.

### Cluster detection and enrichment analysis

Single-molecule trajectories of SctQ, SctL, and SctN recorded under secretion-ON conditions were analyzed to test whether long-lived binding events were spatially enriched. Approved trajectories were converted into individual binding events, each represented by the mean localization position and event duration. Events lasting ≥ 30 frames were classified as long.

Spatial event clusters were identified using DBSCAN (ε = 100 nm, minPts = 2). For each field of view, the observed fraction of long events within each cluster was compared with the global fraction of long events using a one-sided hypergeometric test. Resulting p-values were corrected for multiple testing by the Benjamini–Hochberg procedure (α = 0.05), and significant clusters were optionally validated by 2,000 random label-shuffle permutations. Clusters were categorized as mixed, pure-long, or pure-short based on the composition of event durations.

### Super-resolution fluorescence imaging & Data Processing

Up to 20,000 frames are collected per field of view with an exposure time of 25 ms. Phase contrast images were taken before and after fluorescence imaging. Raw data were processed in MATLAB using a modified version of easy-DHPSF software^35,60^. Fluorescent fiducial markers were used for sample drift correction. Phase contrast images were segmented using Omnipose^61^. Single-molecule localizations were registered to phase contrast masks using a two-step affine transformation in MATLAB, as described previously^35^. Localizations outside of cells were discarded from further analysis using axial bounds and Omnipose-derived cell outlines.

### MSD / cumulative displacement tracking

To derive single-molecule displacements, localizations in subsequent frames were linked into trajectories. A maximum linking distance of 2.5 μm was used for linking analysis, and multiple localizations that were present at the same time within a single cell were discarded to prevent misassignment of molecules.

For each trajectory, the cumulative displacement was calculated as the running sum of all stepwise displacements between consecutive localizations,

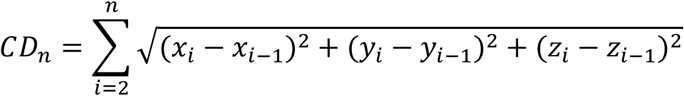

where *x*_*i*_, *y*_*i*_, *z*_*i*_ are the 3D coordinates of the molecule at frame *i*. The cumulative displacement versus time relationship was used to visualize overall molecular motion and to compute the mean step size or slope of displacement accumulation over time. This slope provided a direct, model-free metric of apparent mobility for each trajectory. The plotted mean displacement normalized by time is thus an average of this displacement, which is equivalent to the slope of the cumulative displacement curve over time.

### Estimating injectisome-bound fraction

To identify the fraction of molecules bound, two parallel approaches were taken: RMS based classification of the trajectories, and clustering of the localizations to inform classifications of cluster-associated trajectories and non-associated trajectories. For RMS-based classification, trajectories were taken and the root mean squared radius was plotted. We recognized that injectisome associated trajectories display a very confined radius compared to cytosolic confusion (see Fig S4d). Trajectories could just be classified based on a threshold. For cluster-based classification, all localizations within one cell were taken and a clustering algorithm was applied. To avoid overcounting repeat localizations of the same molecule, trajectories were sorted based on association with a cluster or not.

#### Spatial confinement–based classification of injectosome-bound trajectories

Single-molecule trajectories were reconstructed as time-ordered 3D coordinates (*x*_*i*_, *y*_*i*_, *z*_*i*_). For each trajectory, spatial confinement was quantified as the radius of gyration (RMS radius), defined as

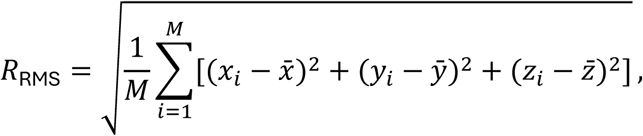

where *M* is the number of localizations in the trajectory and *x^-^, y^-^, z^-^* is the centroid of that trajectory. Trajectories were classified as “injectosome-bound” if *R*_RMS_ < 120nm, and as “cytosolic/mobile” otherwise. For each cell, the fraction of injectosome-bound trajectories was calculated and the percent bound value was reported as bound fraction representing that cell’s injectosome-association (table S5).

#### Detection of subcellular localization clusters

All 3D localizations (*x*, *y*, *z*) within a cell were clustered using density-based spatial clustering (DBSCAN). Two parameters were defined per cell: (i) a spatial neighborhood radius of 60 nm and (ii) a minimum cluster size (minPts). The minimum cluster size was scaled with total localization count in that cell as minPts = round(*N*/300), where *N* is the number of localizations in that cell, with a lower bound of 6 points. Cells for which this procedure would have produced minPts ≤ 3 were excluded from cluster analysis.

Localizations that did not meet these density criteria were labeled as noise and not assigned to a cluster. For each remaining cluster, the centroid was defined as the mean 3D position of all localizations assigned to that cluster.

To determine whether a trajectory was spatially associated with an injectosome, each trajectory from a given cell was compared to that cell’s cluster centroids. For every localization in a trajectory, the Euclidean distance to each centroid was computed, and the minimum distance to the nearest centroid was retained.

A trajectory was classified as “injectosome-bound by proximity” if at least 50% of its localizations lay within 100 nm of any cluster centroid. All other trajectories were classified as unbound.

For every cell we therefore obtained: (i) the number of clusters detected, (ii) the percent bound by cluster proximity, and (iii) the percent bound by RMS confinement. These per-cell values were saved and then aggregated across fields of view, proteins, and conditions for statistical comparisons (table S6).

## Notes

### Competing Interest Statement

The authors have declared no competing interest.

### Summary of Updates

Figure 5 revised; additional supplemental analysis

