## Supplemental information for "Cooperative Binding of Cytosolic Type III Secretion System Proteins to the Injectisome Revealed by Live-Cell Single-Molecule Localization Microscopy"

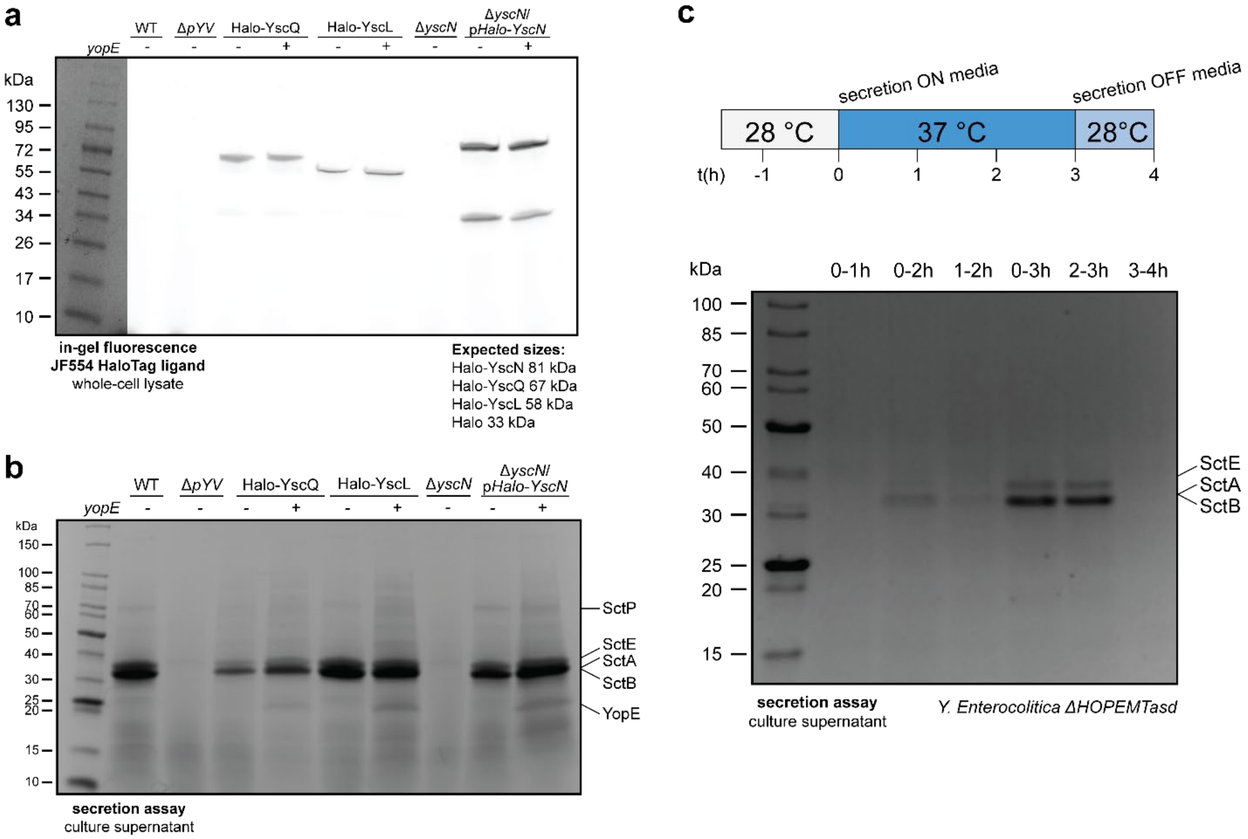

**Figure S1. Validation of Halo-tagged constructs and secretion activity. (a)** In-gel fluorescence confirms expression of Halo-tagged SctQ, SctL, and SctN constructs. Minor cleavage products are visible, particularly for Halo-SctN, consistent with a free HaloTag. For the measurements in this manuscript, a freely diffusing HaloTag would blur into the background rather than contribute to bound measurements (see Fig 1.)
**(b)** Secretion assay confirms that Halo-tagged constructs are secretion-active.
**(c)** Secretion is reversibly induced by calcium depletion and restored by calcium repletion, validating the experimental toggling of secretion ON/OFF states.

### Photobleaching rate of JFX-554 under long exposure imaging conditions

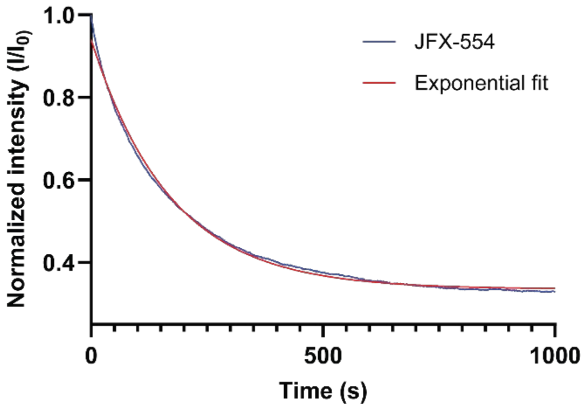

**Figure S2. Photobleaching time constant of JFX-554 under imaging conditions.** Mono-exponential fit of fluorescence decay yields a bleaching time constant of $t_{bleach}$ = 171.5 seconds, confirming suitability of JFX-554 as a label for long-exposure time single-molecule imaging.

To quantify the photobleaching time constant, Halo-SctQ cells were prepared as described (see "Methods - Cell Plating for Microscopy") but stained with 50 nM JFX-554 HaloTag ligand dye. The same long-exposure imaging conditions were used as described in Methods - Optical Setup. The image stack was segmented using Omnipose[1], and the mean fluorescence intensity within each cell was recorded per frame. A background ROI was selected in a cell-free region of the same field of view, and its mean intensity was subtracted from the cellular ROI intensity at each time point to correct for camera offset and nonspecific background signal.

The resulting background-corrected intensity trace I(t) was normalized to its initial value $I_0$ (first frame) to allow comparison across cells and replicates. The normalized decay curve was then fit to a single-exponential decay model,

$$I(t) = A\, e^{-kt}$$

where *A* is the initial normalized intensity (approximately 1 at t = 0) and *k* is the first-order photobleaching time constant. Fits were performed by nonlinear least-squares regression in Graphpad Prism and the best-fit *k* was determined to be 171.5 seconds.

#### Total analyzed events for bound time calculations

| | | Total number of counts |
|---|---|---|
| **Δeffector/ON** | **SctQ** | 1417 |
| | **SctL** | 1463 |
| | **SctN** | 5857 |
| **Δeffector/OFF** | **SctQ** | 4063 |
| | **SctL** | 1973 |
| | **SctN** | 5545 |
| **YopE/ON** | **SctQ** | 1003 |
| | **SctL** | 1902 |
| | **SctN** | 1240 |
| **YopE/OFF** | **SctQ** | 1345 |
| | **SctL** | 1446 |
| | **SctN** | 2292 |

**Table S1. Total number of bound time events recorded for each protein and condition.** Summary of total counts used in bound time analysis for SctQ, SctL, and SctN across four experimental conditions: Δeffector/OFF, Δeffector/ON, YopE/OFF, and YopE/ON.

### Spatial distribution of long- and short- bound modes for T3SS$^{\Delta effector/ON}$

| | Δeffector/ON | | |
|---|---|---|---|
| | SctQ | SctL | SctN |
| **Total number of counts** | 1417 | 1463 | 5857 |
| ***Breakdown of long bound-times (>15 seconds) vs short (<15 seconds)*** | | | |
| **Single-binding events / total** | 288/1417 | 361/1463 | 720/5857 |
| **Multi-binding events / total** | 1129/1417 | 1102/1463 | 5137/5857 |
| Mixed | 112/1129 (9.9%) | 331/1102 (30.0%) | 1766/5137 (34.4%) |
| Pure-short | 1005/1129 (89.0%) | 751/1102 (68.1%) | 3346/5137 (65.1%) |
| Pure-long | 12/1129 (1.1%) | 20/1102 (1.8%) | 25/5137 (0.5%) |

**Table S2. Spatial co-occurrence of short- and long-lived binding events for T3SS$^{\Delta effector/ON}$ conditions.** Quantitative analysis of whether long-bound events occur at the same spatial locations as short-bound events. The observed event-frequencies support the conclusion that individual injectisomes can engage in both binding modes.

### Spatial distribution of long and short binding modes for $T3SS^{YopE/ON}$

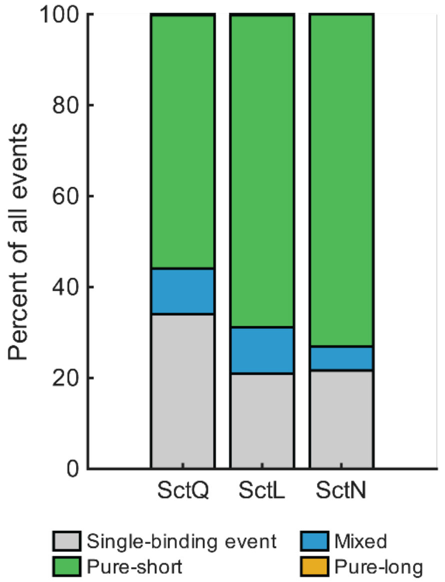

**Figure S3. Bound time analysis of SctQ, SctL, and SctN under YopE/ON conditions.** Bound time distributions for each protein under active secretion in the presence of YopE. All proteins show that within clusters with multiple binding events, there is a mixture of short, long, or mixed bound times. Specific counts for each condition are reported in table S3.

| | | **YopE/ON** | |
|---|---|---|---|
| | **SctQ** | **SctL** | **SctN** |
| **Total number of counts** | 1003 | 1902 | 1240 |
| ***Breakdown of long bound-times (>15 seconds) vs short (<15 seconds)*** | | | |
| **Single-binding events / total** | 341/1003 | 398/1902 | 268/1240 |
| **Multi-binding events / total** | 662/1003 | 1504/1902 | 972/1240 |
| Mixed | 101/662 (15.3%) | 194/1504 (12.9%) | 65/972 (6.7%) |
| Pure-short | 559/662 (84.4%) | 1306/1504 (86.8%) | 907/972 (93.3%) |
| Pure-long | 2/662 (0.3%) | 4/1504 (0.3%) | 0/972 (0.0%) |

**Table S3. Spatial co-occurrence of short- and long-lived binding events for $T3SS^{YopE/ON}$ conditions.** Quantitative analysis of whether long-lived binding events occur at the same spatial locations as short-lived events as visualized in figure S3. The observed event-frequencies support the conclusion that individual injectisomes can engage in both binding modes.

### Single Molecule Localization and Tracking Microscopy of SctQ

| Halo-SctQ single molecule tracking | | |
|---|---|---|
| | **Number of cells** | **Number of trajectories** |
| Δeffector/ON | 193 | 19439 |
| YopE/ON | 355 | 56580 |
| Δeffector/OFF | 381 | 28772 |
| YopE/OFF | 380 | 55397 |

**Table S4. Summary of single-molecule tracking data for Halo-SctQ.** Number of cells and total trajectories recorded for each condition. For each condition, data was collected on at least two different days, from unique cultures.

### Statistical Testing Similarity Across Conditions in Cluster- and RMS-based Bound Fraction

#### Unpaired t-test RMS-based bound fraction

| Comparison | t Statistic | Degrees of Freedom | p Value | Interpretation |
|---|---|---|---|---|
| ON: Δeffector vs +YopE | -2.7357 | 3.6861 | 0.057 | n.s. |
| OFF: Δeffector vs +YopE | 1.0686 | 2.8313 | 0.3678 | n.s. |
| Δeffector: ON vs OFF | -2.4808 | 2.782 | 0.0958 | n.s. |
| +YopE: ON vs OFF | 1.6274 | 3.737 | 0.184 | n.s. |

**Table S5. Statistical comparison of RMS-based bound fractions across secretion state and effector conditions show differences are within statistically expected variation.** Mean bound fractions were computed at the biological sample level and shown in Fig 5c and compared between conditions using two-sided unpaired Welch's t tests. For each comparison, the null hypothesis was that the mean per-sample bound fraction was equal between the two conditions. Reported p values reflect the probability of observing a difference in means at least as large as that measured under the null hypothesis. Nonparametric Wilcoxon rank sum tests produced identical conclusions.

#### Unpaired t-test cluster-based bound fraction

| Comparison | t Statistic | Degrees of Freedom | p Value | Interpretation |
|---|---|---|---|---|
| ON: Δeffector vs +YopE | -1.8845 | 3.8344 | 0.1357 | n.s. |
| OFF: Δeffector vs +YopE | 1.4069 | 1.2736 | 0.3548 | n.s. |
| Δeffector: ON vs OFF | 0.422 | 1.6911 | 0.7205 | n.s. |
| +YopE: ON vs OFF | 2.7182 | 3.885 | 0.0548 | n.s. |

**Table S6. Statistical comparison of cluster-based bound fractions across secretion state and effector conditions show differences are within statistically expected variation.** Mean bound fractions were computed at the biological sample level and shown in Fig 5f and compared between conditions using two-sided unpaired Welch's t tests. For each comparison, the null hypothesis was that the mean per-sample bound fraction was equal between the two conditions. Reported p values reflect the probability of observing a difference in means at least as large as that measured under the null hypothesis. Nonparametric Wilcoxon rank sum tests produced identical conclusions.

**How Does $\tau$ Relate to Turnover at All Binding Sites of SctQ?**

Motivated by an interest in comparing our data to previously acquired FRAP data on SctQ, we derive the expected unbinding time for all binding sites, just as one would expect to probe during FRAP. Following this brief derivation, we will also discuss additional factors that convolute the experimentally observed measurement.

**Unbinding time at a single binding site.**

For a single binding site $i$, the dissociation process is modeled as an exponential random variable, where $T_i$ is the time for site $i$ to unbind and $k_{\text{off}}$ is the dissociation rate constant:

$$T_i \sim \text{Exp}(k_{\text{off}}).$$

The survival probability, i.e., the probability that the site remains bound beyond time $t$, is given by

$$P(T_i > t) = e^{-k_{\text{off}} t}.$$

The expected lifetime of a single site (denoted by $\mathbb{E}[T_i]$) is therefore

$$\mathbb{E}[T_i] = \frac{1}{k_{\text{off}}} = \tau.$$

**Unbinding time for multiple independent binding sites.**

To observe the unbinding of multiple binding sites, the exponential decay is sampled for every binding site $i$. We define $N$ as the number of binding sites, and $T_i$ as the time for site $i$ to unbind. The total unbinding time for the full complex, $T_{\text{full}}$, is

$$T_{\text{full}} = \max\{T_1, T_2, \dots, T_N\}.$$

In other words, we ask: how long does it take until all $N$ independent exponential events have occurred at least once? It is then the slowest binding site that determines when the full ring has unbound.

Let the time of the first unbinding event $T_{(1)}$ be

$$T_{(1)} = \min\{T_1, \dots, T_N\}.$$

All these $T_i$ are independent events, therefore the probability that the first unbinding has not yet occurred by time $t$ is equal to the probability that every binding site is still bound at time $t$, this can be denoted as:

$$P(T_{(1)} > t) = P(T_1 > t,\ T_2 > t, \dots, T_N > t).$$

As a result of this independence, the probabilities factor, thus

$$P(T_1 > t,\, T_2 > t, \dots, T_N > t) = \prod_{i=1}^{N} P(T_i > t).$$

Therefore, the survival probability of the first unbinding event is

$$P(T_{(1)} > t) = (e^{-k_{\text{off}} t})^N = e^{-N k_{\text{off}} t},$$

which has the form of an exponential distribution with rate $N k_{\text{off}}$. Thus,

$$T_{(1)} \sim \text{Exp}(N k_{\text{off}}).$$

So the first unbinding event occurs *N* times faster on average than a single-site unbinding. This leads directly to the expected time:

$$\mathbb{E}[T_{(1)}] = \frac{1}{N k_{\text{off}}}.$$

After this event, $N - 1$ sites remain bound. The associated expected time becomes:

$$\mathbb{E}[T_{(2)}] = \frac{1}{(N-1)\, k_{\text{off}}},$$

$$\mathbb{E}[T_{(3)}] = \frac{1}{(N-2)\, k_{\text{off}}}.$$

Note that subsequent unbinding events refer to unbinding at any binding site $i$, not necessarily adjacent positions. These are also conditional: the expected time starts when the previous unbinding has taken place. It follows that the incremental expected times between successive unbinding events increase as fewer sites remain bound. These concepts are summarized in the diagram below.

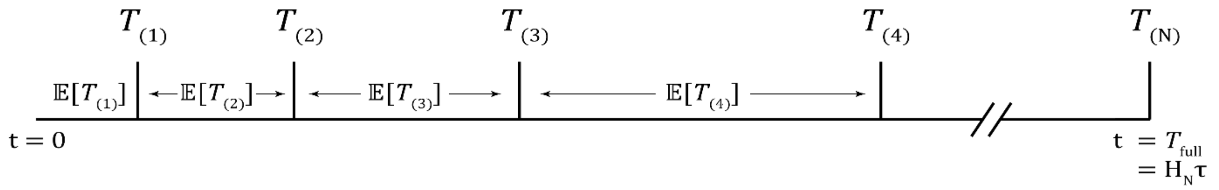

Then the expected time for a full ring of $N$ independent exponential events (each with rate $k_{\text{off}}$) to all occur at least once is therefore, the sum of the above expected times:

$$T_{\text{full}} = \frac{1}{k_{\text{off}}}\left(\frac{1}{N} + \frac{1}{N-1} + \cdots + \frac{1}{1}\right).$$

The sum in parentheses is the $N$-th harmonic number, denoted $H_N$, so the expression can be rewritten as

$$T_{\text{full}} = \frac{H_N}{k_{\text{off}}}.$$

We define the mean lifetime of a single site as

$$\tau = \frac{1}{k_{\text{off}}}.$$

Then the expected time for the full ring to unbind at least once is

$$T_{\text{full}} = \frac{H_N}{k_{\text{off}}} = H_N \tau.$$

**Unbinding time with two distinct lifetimes.**

If a biological system is known to be a mixture of two distinct exponential processes, we can update the above statement. Let the short and long lifetimes be

$$\tau_1 = \frac{1}{k_{\text{off},1}}, \qquad \tau_2 = \frac{1}{k_{\text{off},2}},$$

with corresponding rates $k_{\text{off},1}$ and $k_{\text{off},2}$. Assume that each site independently samples the short lifetime with probability $p$ and the long lifetime with probability $1 - p$.

Thus, each site's dissociation time is a mixture random variable

$$T_i \sim p\,\text{Exp}(k_{\text{off},1}) + (1 - p)\,\text{Exp}(k_{\text{off},2}).$$

The survival function for a single site is

$$P(T_i > t) = p\,e^{-k_{\text{off},1} t} + (1 - p)\,e^{-k_{\text{off},2} t}.$$

The expected lifetime of a single site is the weighted sum

$$\mathbb{E}[T_i] = p\,\tau_1 + (1 - p)\,\tau_2.$$

The expected time for all binding sites to unbind becomes

$$T_{\text{full}} = \frac{1}{p\,k_{\text{off},1} + (1 - p)\,k_{\text{off},2}} \left( \frac{1}{N} + \frac{1}{N - 1} + \cdots + \frac{1}{1} \right) = \frac{H_N}{p\,k_{\text{off},1} + (1 - p)\,k_{\text{off},2}}.$$

Equivalently,

$$T_{\text{full}} = H_N (p\,\tau_1 + (1 - p)\,\tau_2).$$

**Turnover for all SctQ binding sites at the injectisome.**

For SctQ under secretion ON conditions, we measure two bound-state lifetimes: a short lifetime $\tau_1 = 4.5$ s and a long lifetime $\tau_2 = 23$ s. The long-lived state occurs with probability $1 - p = 0.06$, so $p = 0.94$ for the short-lived state.

Using the expression

$$T_{\text{full}} = H_N(p\,\tau_1 + (1-p)\,\tau_2),$$

we obtain

$$T_{\text{full}} = H_N(0.94 \times 4.5\text{ s} + 0.06 \times 23\text{ s}) = H_N(5.61\text{ s}).$$

For a ring with $N = 24$ binding sites, the harmonic number is $H_{24} \approx 3.79$, so

$$T_{\text{full}} = 3.79 \times 5.61\text{ s} = 21.26\text{ s}.$$

Thus, given the observed mixture of SctQ lifetimes, the expected time for all 24 sites in the ring to undergo at least one unbinding event is on the order of 21 s. If these binding sites are rapidly reoccupied (a product of the concentration and the $k_{on}$), this time approximates the time of complete turnover (unbinding followed by binding). Ultimately, FRAP measures both events: once all bleached molecules have been replaced by an unbleached molecule (i.e. unbound and bound), full fluorescence recovery is expected. Next, we can ask if we should expect a FRAP recovery of 21 s.

**Experimentally Observing Complete Ring Turnover.**

Traditionally, FRAP may be used for an ensemble measurement of complete turnover for a protein complex. In practice, FRAP recovery reflects a convolution of binding kinetics, diffusion, and photophysics, and several effects can make the observed recovery time longer than the underlying turnover time, or even prevent complete recovery to the pre bleach level, most prominently:

- **Replacement with bleached subunits.** If the cytosolic pool of proteins is partially bleached during the initial bleach, newly arriving molecules may already be non-fluorescent. Molecular turnover still occurs, but because the incoming subunits contribute no additional new fluorescence, the observed recovery is flattened or delayed relative to the true exchange kinetics.
- **Photobleaching during recovery.** Continuous imaging after the initial bleach unavoidably bleaches additional fluorophores within the field of view. This ongoing loss of fluorescence lowers the apparent recovery plateau and can make recovery appear slower or incomplete, even when every binding site has exchanged because newly arriving molecules may be bleached.
- **Photophysical effects.** Fluorophores can transition temporarily into dark or non-emissive states, resulting in fluorophore blinking. These processes alter the apparent fluorescence independently of binding or unbinding, adding noise or bias to the recovery trace.

Taken together, these effects mean that the theoretically expected ring turnover time of $\sim$ 22 s provides a lower bound on the timescale for complete exchange of SctQ at the injectisome. FRAP experiments may report a longer or incomplete recovery, even when all binding sites have turned over at least once on the microscopic level.

**Observed Bound-State Lifetimes Are Sufficient to Sustain Reported Secretion Rates**

The bound-state lifetimes measured provide a kinetic measure of how long individual proteins remain bound to the injectisome. It has previously been postulated that chaperones handover effectors in the cytosol to SctQ-containing complexes which in turn "shuttle" effectors to the injectisome, driving secretion[2, 3]. Whether these dynamics can sustain previously reported secretion rates[4] (7–60 effector proteins $s^{-1}$ $cell^{-1}$) then depends on how rapidly molecules unbind so they can be reoccupied. In the maximum case, where $k_{on}$ is relatively high and binding sites are saturated, such that each unbinding event is promptly followed by another binding event, then the unbinding rate of bound SctQ directly defines the maximum secretion throughput.

The absence of substantial fluctuations in bound fraction or appearance rate indicates that the number of active binding sites and their occupancy are largely constant, consistent with a saturated regime in which injectisome binding sites are rarely unoccupied. We therefore assume that the unbinding rates of SctQ, L, and N are much slower than the effective binding rate. Under this assumption, the dissociation rate constant provides a direct estimate of the maximum turnover per site. Using the experimentally measured bound-state lifetime of Halo–SctQ under $T3SS^{YopE/ON}$ conditions ($\tau$ = 4.5 s [3.9, 4.9], and 23 s [13, 44]) and the average number of injectisomes per cell (≈ 12) (fig. 5e), we modeled the maximum theorical secretion rate using $\tau_{short}$ (Fig S5). An additional consideration is the number of SctQ copies per injectisome, i.e. the binding sites for SctQ:effector complexes that can be converted into secretion events. The disagreement between available structural density and live-cell measurements motivates our three modeled curves. *In situ* cryo-ET studies in Shigella and Salmonella have consistently resolved six cytoplasmic "pods" arranged around the ATPase[5, 6], implying 6- or 12-copy possibilities if one or two SctQ subunits occupy each pod[7, 8]; other studies[9-12] estimate the SctQ copy number at the injectisome at 24.

Across these different copy number scenarios, the model achieves per-cell secretion rates of approximately 16 – 64 molecules $s^{-1}$ (Fig. S5). This is in agreement with reported values[4] which yield per-cell secretion rates of 7–60 molecules $s^{-1}$. These findings indicate that, under near-saturated conditions, SctQ turnover kinetics could be sufficient to sustain physiological secretion throughput. Under the assumption of 24 binding sites, the other two curves in our model naturally acquire additional meanings: the 12-site curve can be interpreted as either half the physical sites or, equivalently, 50% conversion efficiency/occupancy (i.e., one secretion per two SctQ binding events), and the 6-site curve as either a 6-copy architecture or 25% conversion efficiency/occupancy (one secretion per four binding events). These alternative readings allow the same modeling framework to capture uncertainty in stoichiometry and potential inefficiency in the coupling between SctQ:effector binding and it's conversion to secretion. Collectively, these analyses demonstrate that SctQ turnover at the injectisome occurs on a timescale compatible with the physiological secretion rates.

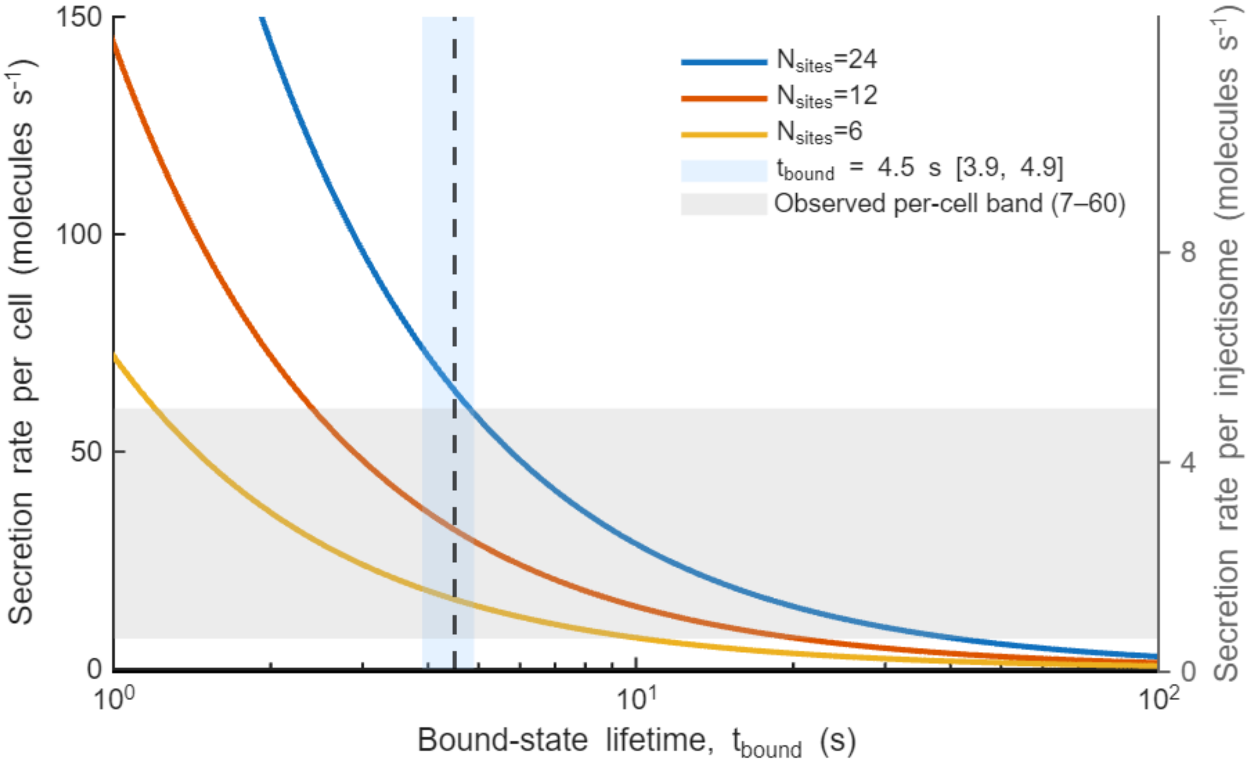

**Figure S5. Predicted secretion rate as a function of SctQ bound-state lifetime.**
Model of secretion output assuming that each SctQ binding event corresponds to one secretion event. The model incorporates the experimentally measured bound-state lifetime of SctQ under secretion ON conditions ($\tau$ = 4.5 s [3.9, 4.9]) and the average number of injectisomes per cell (≈ 12). Curves represent different assumptions about the number of available binding sites per injectisome and the conversion between binding and secretion events: (i, blue) 24 subunits, corresponding to perfect occupancy (1:1 binding = secretion), (ii, orange) 12 subunits, equivalent to half occupancy or half conversion efficiency (1:2 binding = secretion), and (iii, yellow) 6 subunits, equivalent to quarter occupancy or quarter conversion efficiency (1:4 binding = secretion). Across these scenarios, predicted secretion rates (7–60 molecules $s^{-1}$ $cell^{-1}$) overlap with reported secretion rates for *Yersinia enterocolitica*, indicating that the measured SctQ turnover kinetics are sufficient to sustain the experimentally observed secretion flux.
